# Type I-E CRISPR-Cas system as an immune system in a eukaryote

**DOI:** 10.1101/357301

**Authors:** Devashish Rath, Lina Amlinger, Gargi Bindal, Magnus Lundgren

## Abstract

Defense against viruses and other mobile genetic elements (MGEs) is important in many organisms. The CRISPR-Cas systems found in bacteria and archaea constitute adaptive immune systems that acquire the ability to recognize MGEs by introducing nucleic acid samples, spacers, in the CRISPR locus. The CRISPR is transcribed and processed, and the produced CRISPR RNAs guide Cas proteins to degrade matching nucleic acid sequences. No CRISPR-Cas system is found to occur naturally in eukaryotic cells but here we demonstrate interference by type I-E CRISPR-Cas system from *Escherichia coli* introduced in *Saccharomyces cerevisiae*. The designed CRISPR arrays are properly expressed and processed in *S. cerevisiae*. Targeted plasmids display reduced transformation efficiency, indicative of DNA cleavage. Unlike *e.g.* Cas9-based systems, which can be used to inactivate MGEs in eukaryotes by introducing specific mutations, type I-E systems processively degrade the target. The type I-E system thus allows for defense without knowledge of MGE gene function. The reconstituted CRISPR-Cas system in *S. cerevisiae* can also function as a basic research platform for testing the role of various factors in the interference process.

## Introduction

Viruses and other mobile genetic elements (MGEs) are potential threats to most studied cellular organisms, by acting as predators or by reducing fitness. In response, organisms have evolved multiple defense strategies, largely grouped into innate and adaptive systems. Innate systems are characterized by being activated by certain preset features of infection. Adaptive systems, on the other hand, can learn to recognize previously unrecognized pathogens. For a long time the vertebrate adaptive immune system was the only known example, but the CRISPR-Cas systems of archaea and bacteria have been demonstrated to be a *bona fide* adaptive immune systems (1). All studied CRISPR-Cas systems are based on short DNA or RNA sequences (protospacers) from *e.g.* virus genomes being stored as DNA spacers in the Clustered Regularly Interspaced Short Palindromic Repeats (CRISPR) locus. A long precursor CRISPR transcript (pre-crRNA) is processed into CRISPR RNA (crRNA) and used by CRISPR associated (Cas) protein effectors to locate and destroy matching targets. The target can be DNA or RNA depending on the type of CRISPR-Cas system (2, 3). CRISPR-Cas systems are grouped into class 1 and 2, depending on if the effector is a protein complex or a single enzyme, respectively. Each class contains several functionally different types of systems (4).

Programmable nucleases such as Zinc finger nucleases (ZFNs), transcription activator like effector nucleases (TALENs), and Cas9 can function as anti-MGE systems in eukaryotic cells by inducing crippling mutations. Particularly Cas9, which has revolutionized gene editing in eukaryotes, has been demonstrated to effectively target several human viruses (5). In basic Cas9 technology DNA cleavage is directed by a single guide RNA (sgRNA). The break can be repaired by Homology Directed Repair (HDR) or error-prone Non-Homologous End Joining (NHEJ) (2). In the case of viruses, if a gene is repaired by the error-prone NHEJ it may be inactivated, resulting in inability of the virus to proliferate, as demonstrated for *e.g.* HIV1 (6). However, this approach requires thorough understanding of the virus’ biology, as randomly targeting a virus gene does not guarantee reduced proliferation.

In this study we take advantage of the properties of the type I-E CRISPR-Cas system to test its ability to function as an anti-MGE system in eukaryotes that do not require understanding of target gene function. The key factors in target degradation by type I-E systems are the Cascade protein-RNA complex and the Cas3 enzyme. Cascade processes and retains crRNA, and uses the crRNA to identify DNA targets flanked by a protospacer adjacent motif (PAM) (7). Once a target is identified, the Cas3 enzyme is recruited to degrade the target. Unlike the DNA-targeting class 2 enzymes, which perform blunt or staggered double-stranded cut, Cas3 destroys the target in a processive manner (8). This property makes Cascade-Cas3 a poor choice for gene editing but highly suitable for removing virus genomes as repair that restore virus function is less likely to occur after processive degradation. As the effect should be the same irrespective of target sequence, Cascade-Cas3 would be especially advantageous for use against poorly characterized viruses. A type I-E system in a eukaryote could also be beneficial for basic research. It allows testing the role and effect of Cas and non-Cas proteins, as well as other factors, for their effect on CRISPR-Cas immunity, independent of their native cellular background.

As a proof-of-concept we adapted the type I-E system from *E. coli* for use as a programmable anti-MGE system in *S. cerevisiae*, chosen for its role as a eukaryotic model system. As targets we use plasmids, which allow comparison with similar experiments in bacterial systems (9–12).

## Results

### Design and reconstitution of type I-E CRISPR-Cas system in *S. cerevisiae*

Our basic system for expressing Cascade, Cas3, and crRNA in *S. cerevisiae* was different plasmids from the pRSGal series, where each produced one of the components (fig. 1A), but we also designed several alternative systems (fig. 1B–D). pCascade, expressing Cascade proteins, had three cassettes with different *cas* genes under control of bidirectional Gal promoters. The cassettes were separated by *S. cerevisiae* CYC1 terminator to prevent formation of antisense transcripts. For production of targeting crRNA, pCRISPR was constructed by inserting a CRISPR array containing four copies of the J3 spacer (4×J3) (13) under control of a Gal promoter, with a CYC1 terminator preventing read-through. For expression of the Cas3 nuclease we designed pCas3 by cloning *E. coli cas3* under a Gal promoter. An alternative version of the plasmid, pCas3::Cse1, carried a naturally occurring *cas3-cse1* fusion (8). Co-expression of this plasmid with pCascade would result in incorporation of the Cas3-Cse1 fusion in a part of the Cascade complexes. To test the effect of different crRNAs we combined a CRISPR with two alternative spacers, J1 and J2, with Cas3 or Cas3-Cse1 fusion to generate pCRISPR+Cas3 and pCRISPR+Cas3::Cse1, respectively.

**Figure 1.**
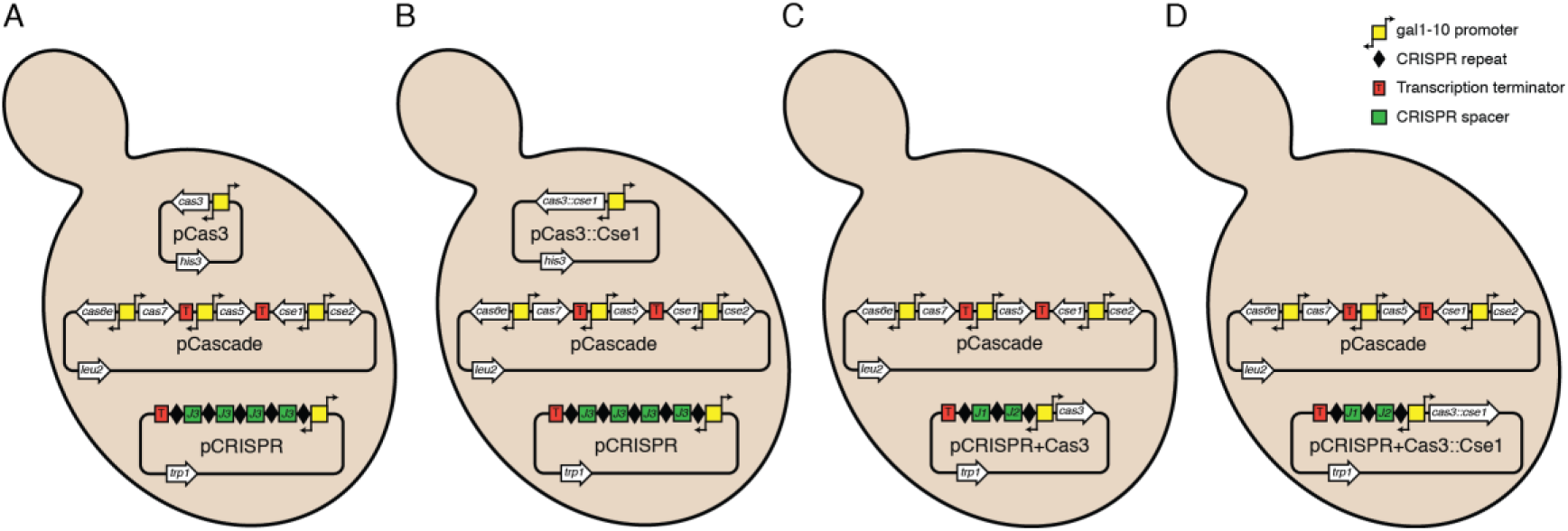
A-D: The different variants of the type I-E CRISPR-Cas system used for expression in *S. cerevisiae*.

Two different target plasmids were designed. pTargetHigh carried a phage Lambda J gene fragment containing the PAMs and protospacers for J1, J2, and J3 (table S4) on a high copy vector. pTargetLow, carried the same J fragment on a low copy backbone. Empty vectors lacking the J fragment were used as non-targeted controls.

### Processing of crRNA in *S. cerevisiae*

To test function of crRNA processing in yeast we analyzed the formation of crRNA in *S. cerevisiae* W303 after expression from pCascade, pCas3, and pCRISPR was induced by addition of galactose. The analysis was performed by northern blot with a radioactively labeled probe complementary to the J3 spacer. The assay demonstrates formation of a 61 nt RNA species, equal in size to crRNA produced in *Escherichia coli*. The amount of crRNA produced was assayed two, four, and five hours after induction, and crRNA was detected at all time points (fig. 2A).

**Figure 2.**
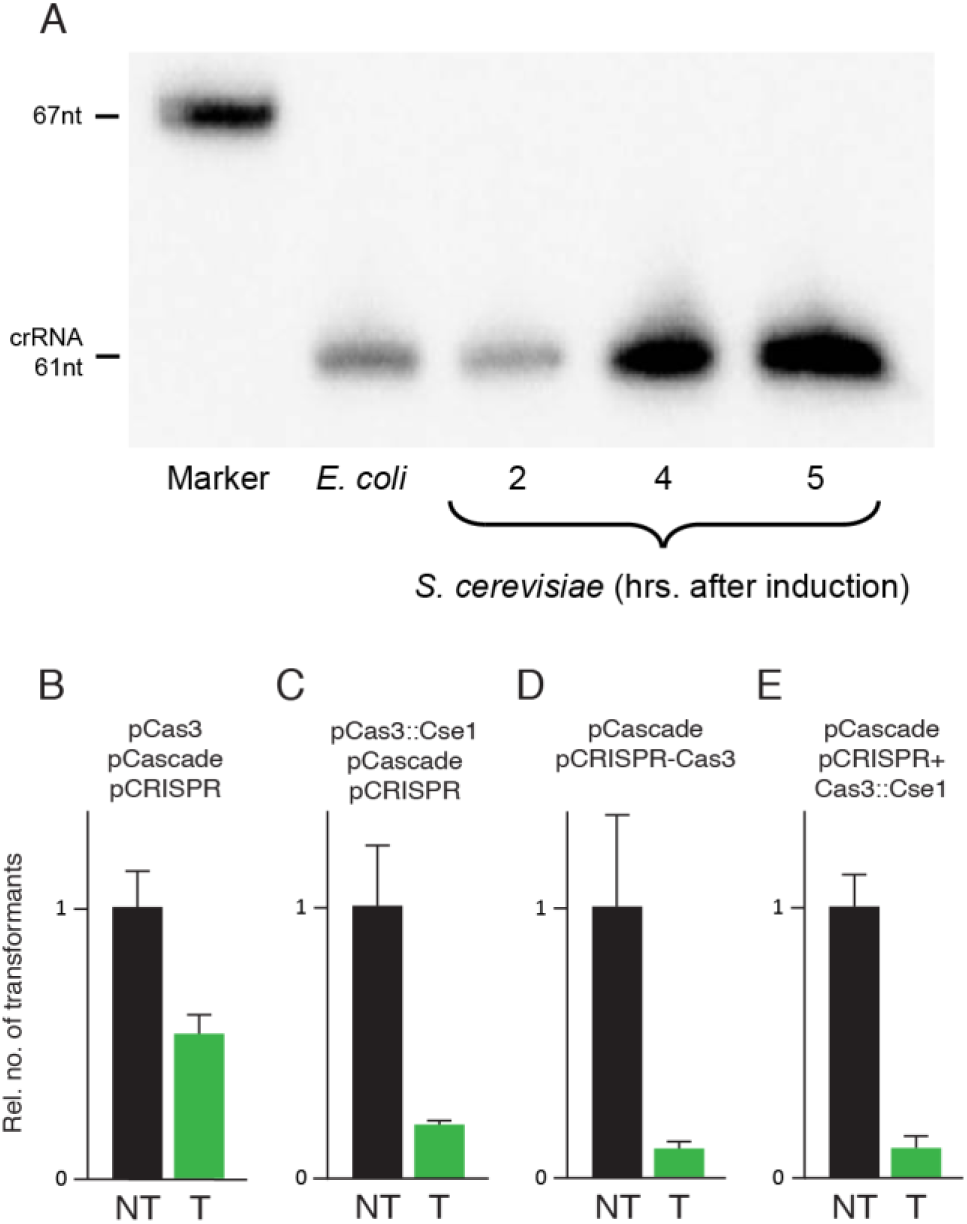
A: Analysis of crRNA by northern blot. *S. cerevisiae* samples were taken at indicated time points after induction of *cas* genes and 4xJ3 CRISPR array expression. RNA from an *E. coli* strain active for interference was used as positive control. B-E: Analysis of CRISPR-Cas interference on plasmids in different *S. cerevisiae* strains (A-B: W303, C-D: BY418) with indicated CRISPR-Cas expression system. Transformation efficiency was measured for plasmids containing a target sequence (T) and non-targeted parent plasmids (NT). Data in B-E is an average of three independent biological replicates normalized so that relative level of transformation by the non-target plasmid is equal in the different panels. Error bars indicate one standard deviation.

### Interference by CRISPR-Cas in *S. cerevisiae*

Functionality of Cascade-Cas3 interference was assessed by comparing transformation efficiency (TE) of a plasmid targeted by the CRISPR-Cas system with that of the same plasmid lacking the target fragment. Transformations were routinely done 240 min after induction of *cas* and CRISPR expression. Using pCas3, pCascade, and pCRISPR in *S. cerevisiae* W303, a 47% decrease in TE of pTargetHigh was observed (fig. 2B). Tests with 150 min induction resulted in similar results (fig. S1). Alternative systems for expressing Cascade, Cas3, and crRNA were tested for improved target interference. Cas3-Cse1 fusion construct, pCas3::Cse1, co-expressed with pCascade and pCRISPR resulted in approximately 73.5% reduction in TE of pTargetHigh in *S. cerevisiae* W303 (fig. 2C). Expression of pCRISPR+Cas3 or pCRISPR+Cas3::Cse1 together with pCascade in *S. cerevisiae* BY418 resulted in 88.6 % and 88.7 % reduction in TE of pTargetLow, respectively (fig. 2D and 2E).

### Effect of interference

Interference assays demonstrated that CRISPR-Cas targeting reduced TE but transformants were still observed on selective plates. These transformants could be carrying target plasmids that had not been cleaved by Cascade-Cas3, plasmids that had been cleaved but subsequently been repaired or plasmids that had acquired escape mutations in the region targeted by CRISPR. NHEJ repair or presence of different escape mutations would result in target sequence heterogeneity in the population of transformed plasmids. To determine sequence heterogeneity in the transformed plasmids we performed the Surveyor assay, which cleave DNA with mismatched bases formed by denaturing and reannealing DNA strands with sequence differences. The target region of pTargetHigh was amplified by PCR and analyzed by Surveyor assay. Sequence heterogeneity at J3 target site would result in two cleavage fragments, both of about 160–170 bp. No such cleavage products could be detected in strains expressing pCRISPR, pCascade and pCas3, similar to interference-negative control strains (fig. S2).

## Discussion

We demonstrate successful reconstitution of bacterial type I-E CRISPR-Cas interference in the eukaryote *S. cerevisiae*. We detect accurate processing of crRNA by northern blot and, importantly, a reduction of target plasmid transformation rate compared to a non-targeted control plasmid. We could not detect sequence heterogeneity in plasmids recovered from transformed cells, demonstrating that the level of interference observed was due to the activity of Cascade-Cas3 and not by varying levels of escape mutations in the target. The lack of sequence heterogeneity also demonstrates that NHEJ repair could not be detected, though this type of repair is rare in *S. cerevisiae*. Our results suggest that the programmable processive DNA degradation by type I-E CRISPR-Cas system can be restored in eukaryotic cells.

In our initial design, Cascade, Cas3, and pre-crRNA is expressed from different plasmids, and this resulted in 50% reduction in plasmid transformation rate. To improve activity we developed improved systems. Co-expression of a Cas3-Cse1 fusion results in a subpopulation of Cascade complexes permanently co-localized with Cas3, which reduced transformation by 73.5 %, a level similar to when J3 crRNA is used against phage Lambda in *E. coli* expressing the LeuO activator (13). We hypothesize that this fusion improves the kinetics of target degradation, as the factors do not need to find each other. Further, we altered the CRISPR to produce two different crRNA targeting the plasmid to potentially increase activity and at the same time expressed Cas3 and pre-crRNA from the same plasmid to reduce the load of carrying multiple plasmids. These modifications resulted in further reduction of transformation, and combination of all improvements result in about one-in-ten plasmids escaping degradation.

Different crRNAs result in different levels of CRISPR-Cas system interference (7, 14, 15), so other spacer sequences may further increase efficiency of target degradation in yeast. We also speculate that the Cascade and Cas3 in the absence of nuclear localization signals (NLS) locate to the cytoplasm, thus allowing only a narrow window of opportunity to destroy their target plasmids while they are en route to the nucleus. This likely causes a reduction in target access and interference compared to bacteria and archaea, which typically lack a membrane around their nucleoid. Other factors that may further improve interference in *S. cerevisiae* are increased expression of Cas protein and pre-crRNA, codon optimization, prevention of detrimental post-translational modification of Cas proteins, and co-expression of *E. coli* host factors that may assist interference.

The large type I-E system is more difficult to establish heterologously than the more compact class 2 systems. However, complete target clearance by type I systems is a distinct advantage over inactivation of MGEs by mutagenesis using class 2 systems. To inactivate an MGE by mutation, thorough understanding of the MGEs function is required to find a key component to target. Also, compared to Cas9-based systems, type I systems can easily target several different sequences by adding more spacer-repeat units to the CRISPR (14, 16), while current applications of Cas9 in eukaryotes require insertion of multiple single guide RNA genes to achieve multiplexing (17). Another benefit of our establishment of a type I-E system in *S. cerevisiae* is that it functions as a platform for testing our current understanding of the system in a non-native background. Such a platform would allow introduction and exclusion of *e.g.* various proteins and other factors component to test their role in the immune system. Finally, while we demonstrate interference in a eukaryote, further development could reconstitute the entire adaptive capability of the system. This would require expression of Cas1 and Cas2, and possibly also other factors such as IHF, indicated as important for adaptation (18). As Richard Feynman stated, “what I cannot create I do not understand”.

## Materials and Methods

### Strains and culture media

All cloning work was done in *E. coli* strain Top10. *E. coli* was cultured in Lysogeny Broth (LB) with aeration at 37°C. When necessary, the media was supplemented with kanamycin (50 μg/ml), ampicillin (100 μg/ml), streptomycin (50 μg/ml), tetracycline (25 μg/ml), and chloramphenicol (15 μg/ml). For experiments in yeast *Saccharomyces cerevisiae* W303 and BY418 strains were used. *S. cerevisiae* BY418 have full deletion of chromosomal copies of genes used for selection, unlike W303 where they are inactivated by point mutations. Yeast cells were cultured in SC medium (Yeast nitrogen base 0.34 %, ammonium sulfate 0.5%, raffinose 2%, quadruple SC dropout mix (CSM, -His, -Leu, -Trp, -Ura; Formedium™, UK) at the rate of 1400 mg/L of medium, pH 5.6). For certain induction experiments SC medium with 1% raffinose was used. When necessary, the media was supplemented with adenine sulfate (80 μg/ml), uracil (20 μg/ml), tryptophan (40 μg/ml), histidine (20 μg/ml), and leucine (60 μg/ml). 1.5% agar was included in the medium to prepare solid medium. Yeast strains were grown with aeration at 30°C. 2% galactose was added to the medium for induction of Gal promoter as required. For a complete list of strains used see table S1.

### Construction of vector expressing Cascade complex

A Gal1-10 promoter cassette from pRS425Gal was cloned between EcoRI/BamHI sites of pBlueScript II SK (+) resulting in pUDM101. All *cas* genes were amplified by colony PCR from *E. coli* MG1655, see table S2 for a complete list of primers. The *cse1* and *cse2* genes were serially cloned into the pUDM101 XbaI and EcoRI sites, respectively, to generate pUDM107. Similarly, *cas7* and *cas6e* were serially cloned in pRS425Gal BamHI and SalI sites, respectively, to generate pUDM108. The four genes were combined by subcloning a HindIII-NotI blunt fragment from pUDM107 into the blunted SacI site of pUDM108, thereby generating pUDM109. The *cas5e* gene was cloned into pRS425Gal vector to generate pUDM110. CYC1 terminator cassette was PCR amplified from pYES2 and subcloned into PstI site of pBlueScript II SK (+) to generate pUDM102. A CYC1-*cas5e*-CYC1 cassette was constructed by serially cloning EcoRV-SmaI fragment from pUDM102 into the SalI and SpeI sites respectively of pUDM110 to generate pUDM315. Using XhoI and NotI this cassette was excised from pUDM315, blunted and sub-cloned into the NotI site of pUDM109 to generate the final Cascade-expressing construct, pCascade. See table S3 for complete list of plasmids used in this study.

### Construction of vectors expressing Cas3 or Cas3-Cse1 fusion

The *cas3* gene was PCR amplified from MG1655 to introduce flanking EcoRI sites and cloned into the EcoRI site of pRS423Gal resulting in the Cas3-expressing plasmid pCas3. A Cas3-Cse1 fusion fragment was amplified from pWUR657 and cloned between the SpeI and NotI sites of pRS423Gal to generate pCas3::Cse1 expressing the Cas3-Cse1 fusion.

### Construction of vectors expressing CRISPR RNA

For expression of crRNA, two CRISPR cassettes, one containing 4xJ3 spacer (13) PCR amplified from pWUR630 and the other containing J1-J2 spacers (a synthetic array ordered from ThermoFisher Scientific Inc.) were cloned into BamHI-NotI sites of pRS424Gal_Cyc1 to place CYC1 terminator directly downstream of the CRISPR array. All spacers targeted the J gene from bacteriophage Lambda. The J1-J2 CRISPR, along with the flanking CYC1 terminator, was excised as a BamHI-SacI fragment and cloned in pCas3 cut with the same sites to generate pCRISPR+Cas3. Additionally, the same BamHI-SacI fragment was blunted and cloned in the blunted SalI site of pCas3::Cse1 to generate pCRISPR+Cas3::Cse1. See table S4 for details of CRISPR and targets.

For expression crRNA in *E. coli*, a minimal CRISPR array with 54 nt of the leader, the J3 spacer and two repeats was cloned into pZE12Luc (19) under the control of PLlacO-1. The plasmid was amplified using primers PLlacO-C and pZE-Xba and the minimal CRISPR array was amplified from pWUR564 (13) using primers LA009 and LA013. The array was cloned by blunt-end ligation in the leader-end so that the first position of the partial leader corresponds to transcription start of the PLlacO-1 and in the other end into the XbaI-site of pZE12Luc.

### Construction of *S. cerevisiae* expressing type I-E CRISPR-Cas system

The vectors constructed as described above were transformed into the desired yeast strains as required. Transformation of *S. cerevisiae* was performed using lithium acetate method (20).

### Construction of target vectors

To construct pTargetHigh, the 350 bp region of Lambda J from pWUR610 was cloned into the BamHI and HindIII sites of pRS426Gal. The cloning replaced the Gal-promoter fragment with the lambda DNA fragment. A low copy target vector, pTargetLow, was constructed by PCR amplification of a region of Lambda J gene using pTargetHigh as template with DR0015 and DR0016 primers and cloning the fragment in the NheI site of pPS1739.

### Analysis of crRNA processing

#### Sample preparation and RNA purification: Yeast

*S. cerevisiae* W303 carrying pCascade, pCRISPR (4xJ3 spacer), and pCas3 was grown to stationary phase, diluted in fresh SC medium and grown with aeration at 32°C to an OD600 value of 0.3. Galactose was then added to induce the CRISPR-Cas system. 10 ml culture was harvested by centrifugation 2, 4, and 5 hours after induction. Total RNA was isolation using the hot phenol method (21).

#### Sample preparation and RNA purification: Bacteria

Overnight culture of *E. coli* BL21AI harboring pWUR397 (Cas3), pWUR400 (Cascade) and pLA002 (minimal J3 CRISPR array), was diluted and grown to OD600 ≈ 0.3. Cas-protein and crRNA expression was induced by addition of 0.2 % arabinose and 1 mM IPTG. After 30 min, 5 ml sample was taken and mixed with 1 ml stop solution (5 % phenol, 95 % ethanol), and pelleted at 4 °C. RNA was purified as previously described (22).

#### Gel and blotting

Samples were run on a 10% denaturing polyacrylamide gel (10 % polyacrylamide, 7 M urea, 1X Tris-borate-EDTA (TBE)) in 1X TBE. Samples were mixed with loading dye (95 % deionized formamide, 0.5 mM EDTA, 0.025 % bromphenol blue/xylene cyanol, and 0.025 % SDS), and boiled 3 min prior to loading. The same was done with the pre-labelled pUC8 size marker. 15 μg of total RNA was loaded for each sample. As a positive control for processed crRNA, total RNA purified from the *E. coli* BL21AI carrying pWUR 397, pWUR400 and pLA002, able to prevent phage infection and plasmid transformation (data not shown), was also loaded on the gel. Transfer to Hybond N+ membrane (Amersham) was done at 4°C overnight at 200 mA. The membrane was UV crosslinked and prehydridized in Church buffer (0.25 M sodium phosphate buffer pH 7.2, 1 mM EDTA, and 7% SDS) for 1 h at 42°C after which 0.5 µM of radioactively labelled probe was added to the buffer. Hybridization was done overnight. The membrane was washed two times 5 min with 2x SSC, 0.1% SDS and exposed the PharosFX-system (BioRad).

The pUC8 size marker (Fermentas) was radioactively labeled using γ-^32^P-ATP (PerkinElmer) and PNK (Fermentas) in an exchange reaction according to the manufacturer’s instructions. The probe, LA014, was labelled the same way in a forward reaction. Excess γ-^32^P-ATP was removed using Illustra ProbeQuant G-50 Micro column (GE Healthcare) purification according to the manufacturer’s instructions.

### CRISPR-Cas activity assays

CRISPR-Cas activity was assessed by plasmid interference assays. Yeast cultures grown to stationary phase and diluted in fresh SC medium containing either 1% or 2% raffinose as carbon source and grown with aeration at 32°C to an OD600 value of 0.2, followed by addition of galactose inducer. 150–240 min after induction the cells were harvested by centrifugation. The cells were then transformed with target vectors or non-target control vectors using lithium acetate method (20), but excluding carrier DNA. Plasmid interference by the CRISPR-Cas system was measured by plating cultures transformed with target vector or non-target vector on selective media and comparing TE.

### Surveyor assay

*S. cerevisiae* BY418 cells harboring pCascade, pCRISPR, and pCas3::Cse1, or negative controls lacking pCRISPR, pCascade or both were grown in SC medium with or without the galactose inducer and transformed with pTargetHigh. The transformed cells were grown for 24 hours in SC medium, pelleted by centrifugation and grown again in equal volume of fresh medium for 24 hours.1.5 ml of the culture was pelleted by centrifugation and resuspended in 500 μl of distilled water. The cells were lysed by heating at 98°C for 10 minutes and 0.5 μl of the lysate was used as template for PCR with primers DR003 and DR004 to amplify the target region. The PCR products were analyzed by electrophoresis on 2 % agarose gel and used for mutation detection with Surveyor mutation detection kit (IDT) as per manufacturer’s instructions. The result of Surveyor assay was analyzed by Qiagen QIAxcel advanced gel electrophoresis system.

## Acknowledgements

We are deeply grateful to Erik Johansson (Umeå University), Hans Ronne (Swedish University of Agricultural Sciences), Gerhart Wagner (Uppsala University), Pernilla Bjerling (Uppsala University) and their respective research groups for providing assistance, strains, and plasmids for this project. This project was funded by the Wenner-Gren Foundations, the Swedish Research Council and the Royal Swedish Academy of Sciences.

## Author Contributions

ML conceived the concept. DR, GB, and ML performed experimental design and DR, LA, and GB performed the experiments. DR and ML analyzed the data and wrote the article.

## Conflict of Interest

The authors declare no conflict of interest.

